# Metastasis and cancer stem cell markers identify two colorectal cancer subtypes with different molecular and clinical features

**DOI:** 10.1101/2020.12.21.423844

**Authors:** Burair A. Alsaihati, Shaying Zhao

## Abstract

Colorectal cancer (CRC) is among the top prevalent cancer types with lethal outcome in the United States and worldwide. CRC inter-tumor heterogeneity highlights the importance of identifying molecular markers for meaningful classification and prognosis. The recently published Consensus Molecular Subtypes (CMS) represent a widely used molecular subtyping system of CRC. However, our analyses indicate that clear heterogeneity still exists in some CMSs. In this work, we demonstrate that both CMS2 and CMS4 are composed of two molecularly distinct subtypes. We named them S1 and S2, short for subtype 1 and subtype 2. Our results indicate that the two subtypes also differ clinically. Notably, S2 exhibits more frequent lymphatic invasion across CRCs and more frequent metastasis events within CMS2 patients.

## Introduction

Colorectal cancer (CRC) is among the top prevalent cancer types with lethal outcome in the United States and worldwide [1, 2]. Recent effort to molecularly classify CRC led to the introduction of the Consensus Molecular Subtypes (CMS) by the CRC Subtyping Consortium [3]. In this subtyping system, CRC is classified into four classes: CMS1, CMS2, CMS3, or CMS4. CMS2 is the most common subtype (∼37% of CRC cases), featured by microsatellite stability (MSS), chromosomal instability (CIN) and favorable survival [3]. Despite its favorable outcome, CMS2 poses a notable metastasis risk, having the second highest frequency of Stage IV [3]. This highlights the need to further investigate CMS2 to identify molecular features that promote its spread and lethality.

In this work, we performed unsupervised clustering of hundreds of CRC cases using CRC metastasis and cancer stem cell (CSC) markers. We identified a simple signature that classifies CMS2 cancers into two subtypes: S1 and S2. Further analysis reveals that within CMS2, S2 is associated with the activation of tumor immune and stromal components, as well as increased lymphatic invasion and metastasis events. Our findings will be useful to the prognosis and precision medicine for CRC patients with CMS2.

## Methods

### Datasets

We assembled a set of 19 gene markers from 7 different publications [4-10]. We downloaded the gene expression for 622 CRC samples from the TCGA Data Portal [11, 12]. We retrieved the microarray gene expression for 566 patients and their clinical data from the Gene Expression Omnibus [13, 14], under accession number GSE39582 [15]. We obtained the CMS labels for both datasets from the synapse repository (https://www.synapse.org/#!Synapse:syn4978511) [3]. We obtained the methylation status, mutation densities and copy number alterations for 459 TCGA CRC samples from a published study [16]. We retrieved the clinical data for TCGA samples from Broad GDAC Firehose version 2016_01_28 [17]. Cancer hallmark gene sets were retrieved from MSigDB [18, 19]. We retrieved published data for the stemness index scores (mDNAsi and mRNAsi) of TCGA samples [20] and gene signatures of tumor and immune microenvironments [21-26].

### Microarray gene-probe selection

For the analysis of immune cell signatures, we used published microarray-based signatures represented by probe ids [21]. For tumor microenvironment cell type signatures and cancer hallmarks, we selected the microarray probe with the highest sample average expression for each gene. For the 19 gene marker co-expression analysis and S1/S2 subtyping, we first built 10 linear regression models (5 using normal and 5 using tumor samples). In these models, the *x* values represent the sample medians of log-transformed Fragment Per Kilobase per Million mapped reads (FPKM) values from TCGA whereas the *y* values represent the sample medians of the corresponding probes with the highest average. Each model uses 300 random genes. From 10 models, we selected the model with the highest correlation coefficient for probe selection. The selected model is represented by: probe sample median = 4.82 + 1.28 × median (log[FPKM + 1]). The list of the selected 19 microarray probes is available in the supplementary.

### Correlation and co-expression analysis

In all our analyses of TCGA gene expressions, we used log-transformed FPKM values: log[FPKM + 1]. The microarray dataset is already preprocessed via Robust Multichip Average (RMA) normalization and batch effect removal [15].

We used a Correlation Score for our gene co-expression analysis. To calculate the Correlation Score between two genes across tumor samples, the score is first set to 0. Then, 40 samples are randomly selected, and the Pearson correlation is calculated. If the correlation is significant (*P* ≤ .05), the Correlation Score is altered by one (incremented if the Pearson correlation is positive or decremented if negative). Otherwise, the Correlation Score is not changed. The process is repeated 100 times using 40 random samples each time so that each gene-pair is assigned a Correlation Score between −100 and 100.

For the co-expression analysis, we used the default hierarchical clustering function, *hclust*, in R version 3.5.1 [27]. For clustering stability analysis, we repeated the calculation of the Correlation Score matrix and the hierarchical clusters 100 times. After each run, we divided the clustering tree into four clusters, using R’s *cutree* function (*k*=4), and examined the result for the presence of specific gene clusters.

### Principal component analysis (PCA), PC1/PC2 values and S1/S2 subtypes

Following the gene co-expression analysis, we merged TCGA colon and rectal samples together. We selected 9 gene markers that form stable clusters to subtype TCGA CRC samples, namely: *VEGFC, SMAD4, TNIK, ITGB1, CD44, ALCAM, VEGFB, OTUB1* and *DLL4*. We used their log-transformed FPKM values for the PCA. Before performing the PCA, we removed expression outliers for each gene. For outlier removal, per gene, all samples with expression higher than the 95^th^ percentile were assigned the 95^th^ percentile value. Using the PCA result from TCGA data, we obtained the gene loadings (coefficients) for the first (PC1) and the second (PC2) principal components and used them to calculate the PC1 and PC2 values for the microarray samples. The cutoff values to separate S1, S2 and mixed samples were determined by heatmap visual examination of PC2-sorted samples. The cutoff selection aims to maximize the sample size of S1 and S2 subtypes while preserving their gene expression contrast. Gene loadings, PC1/PC2 values and S1/S2 labels are all available in the supplementary.

### Cluster centroid comparison

We used permutation test to find the significance of the distance between two clusters, C1 and C2, each consisting of multiple points in *M* dimensions. First, we normalized the values of each dimension. Then, we calculated the Euclidian distance between C1 and C2 centroids (*d*_*o*_). We then ran 1000 permutations. In each permutation, we randomized C1 and C2 assignments of the original points, preserving each cluster’s size as is. In each run, we calculated the permutation distance *d*_*p*_ the same way we calculated *d*_*o*_. Let the number of runs where *d*_*p*_ ≥ *d*_*o*_ be *r*. The permutation *P* value is given by *P* = *r*/1000.

### Enrichment Scores

We calculated the Enrichment Scores based on *P* values in all our results: |Enrichment Score| = −log_10_(*P*). We assigned a negative sign to each S1 enrichment to distinguish it from S2 enrichment, and to depletion to distinguish it from enrichment.

For the immune cell and tumor microenvironment Enrichment Scores, we first divided each CMS class into two groups: signature-high, with ssGSEA scores above the median, and signature-low, with ssGSEA scores below the median. We then used Fisher’s exact test to see if either group is enriched in S1 or S2 of any CMS class. Enrichments for signature-high groups were assigned positive Enrichment Scores and enrichments for signature-low groups were assigned negative Enrichment Scores. We compared the overall survival (OS) between the two groups using Kaplan-Meier log rank test.

## Results

### A simple linear model guides microarray probe selection for each of the 19 gene markers

We performed our analyses in this work using two gene expression datasets: TCGA colon and rectal cancers (622 patients) and a microarray CRC dataset (566 patients) as described in our Methods and Table 1. We collected 19 gene markers previously linked to CRC metastasis and CSCs as described in Table 2.

**Table 1:** A description of the gene expression datasets

| **Dataset** | **TCGA COAD & READ** | **GSE39582** |
| --- | --- | --- |
| Organization | NIH | Ligue Nationale Contre le Cancer |
| Platform | Illumina RNA-Seq | Affymatrix GPL570 |
| MSI-H/dMMR | 87 | 75 |
| MSI-L | 101 | NA |
| MSS/pMMR | 429 | 444 |
| CMS1 | 76 | 91 |
| CMS2 | 219 | 232 |
| CMS3 | 72 | 69 |
| CMS4 | 143 | 127 |
| Total tumor | 622 | 566 |
| Total normal | 51 | 19 |

**Table 2:** Colorectal cancer metastasis and CSC markers

| **Name in literature** | **Current gene symbol** | **Marker type** | **Source** |
| --- | --- | --- | --- |
| CD133 | PROM1 | CSC marker | [9] |
| CD29, ITGB1 | ITGB1 | CSC marker and metastasis promoter | [4, 9, 10] |
| CD44 | CD44 | CSC marker | [4, 9] |
| CD326 | EPCAM | CSC marker | [4] |
| Cripto-1, TDGF1 | TDGF1 | CSC marker and metastasis promoter | [4, 8] |
| CD26 | DPP4 | CSC marker | [4] |
| LGR5 | LGR5 | CSC marker and metastasis promoter | [4, 6] |
| DLL4 | DLL4 | CSC marker | [4] |
| CD166 | ALCAM | CSC marker | [4] |
| Ezrin | EZR | metastasis promoter | [5] |
| VEGF | VEGFA | metastasis promoter | [5] |
| VEGF | VEGFB | metastasis promoter | [5] |
| VEGF | VEGFC | metastasis promoter | [5] |
| VEGF | VEGFD | metastasis promoter | [5] |
| SMAD4 | SMAD4 | metastasis suppressor | [5] |
| IMP3 | IGF2BP3 | metastasis promoter | [5] |
| TNIK | TNIK | metastasis promoter | [5] |
| S100A2 | S100A2 | metastasis promoter | [5] |
| OTUB1 | OTUB1 | metastasis promoter | [7] |

Representing gene expression levels using microarray data can be challenging because most microarray libraries use multiple probes per gene. A previous study has recommended representing each gene by the probe with the highest sample average [28]. Because of the small size of our marker gene set, we needed high consistency between TCGA and microarray gene expression data. Therefore, we opted for an approach based on simple linear regression and guided by the TCGA FPKM values to select representative probes (see Methods). We fitted 10 linear regression models. The results indicate a strong correlation between the microarray probes and their corresponding genes’ FPKM values in TCGA (*P*<2.2E-16 and Pearson *R*^2^>0.5; Figure 1). We selected the linear model with the highest *R*^2^. For each of the 19 markers, we selected the probe that best fits this model (Table S1). Most of the selected probes are indeed these with the highest sample average.

**Figure 1.**
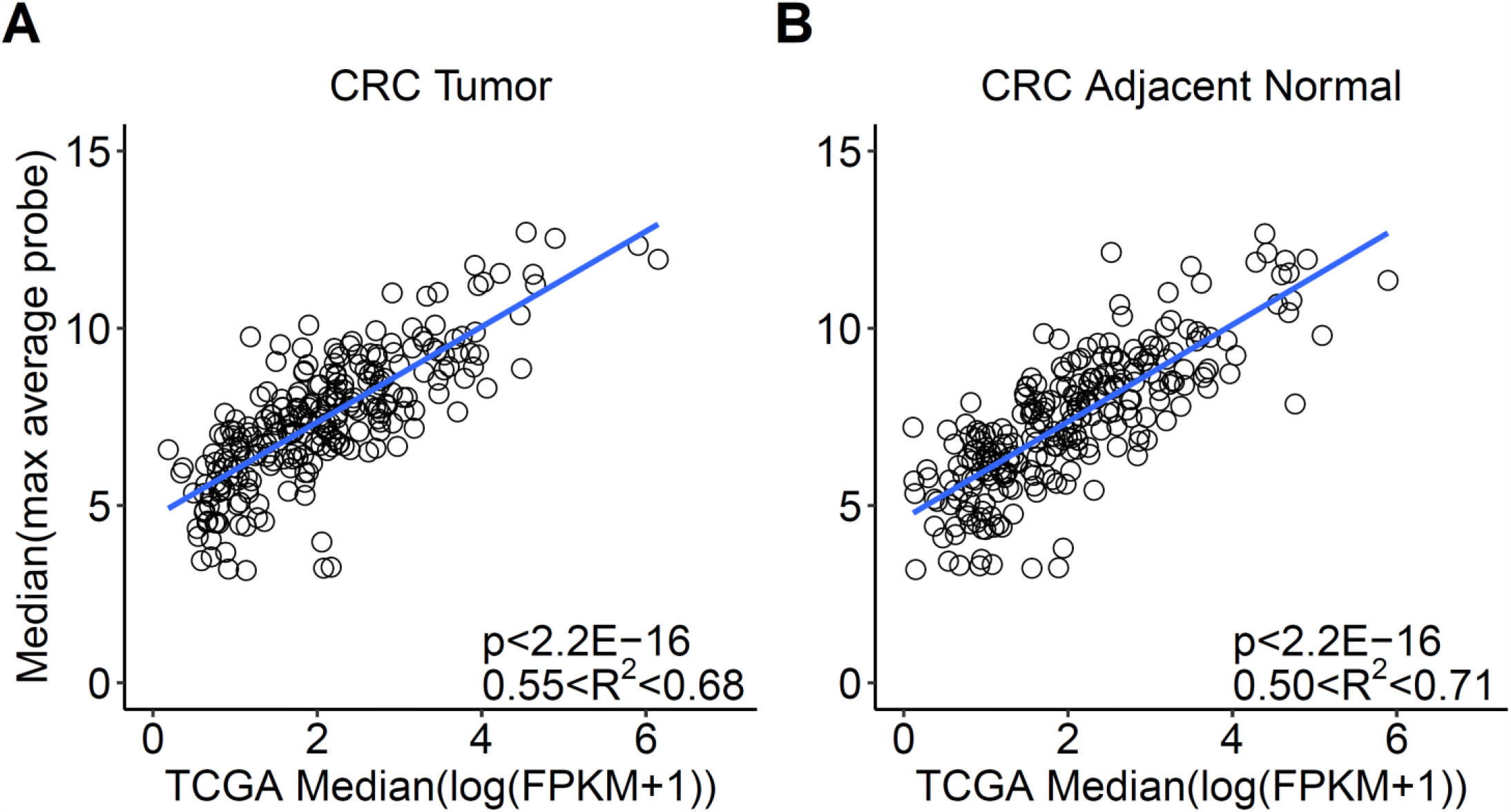
A simple linear model guides microarray probe selection for each of the 19 gene markers. **A**. A representative linear regression model built using CRC tumor samples by fitting the sample median expression values of 300 randomly selected genes. The *x*-axis is the median log transformed FPKM value from TCGA. The *y*-axis is the median expression value for the probe with the highest sample average corresponding to the same gene. Each circle represents one of the 300 genes. Indicated in the bottom right corner, the range of the regression *R*^2^ and the correlation *P* values obtained from five runs, sampling different genes in each run. **B**. A similar representative model built using CRC adjacent normal samples.

### The co-expression pattern of 9 gene markers cannot be explained by the microsatellite (MS) status alone

We defined a Correlation Score value based on random sampling (see Methods) for gene clustering. Our correlation method helps increasing the correlation signal and reducing the effect of gene expression experimental noise. The hierarchical clustering of these Correlation Scores results in two consistent clusters across TCGA colon and rectal samples. Between colon and rectal results, six of 7 genes are shared in Cluster 1: *ALCAM, TNIK, CD44, ITGB1, SMAD4* and *VEGFC*. In Cluster 2, consists of three genes, all genes are shared: *DLL4, VEGFB* and *OTUB1* (Figures 2A and B).

**Figure 2.**
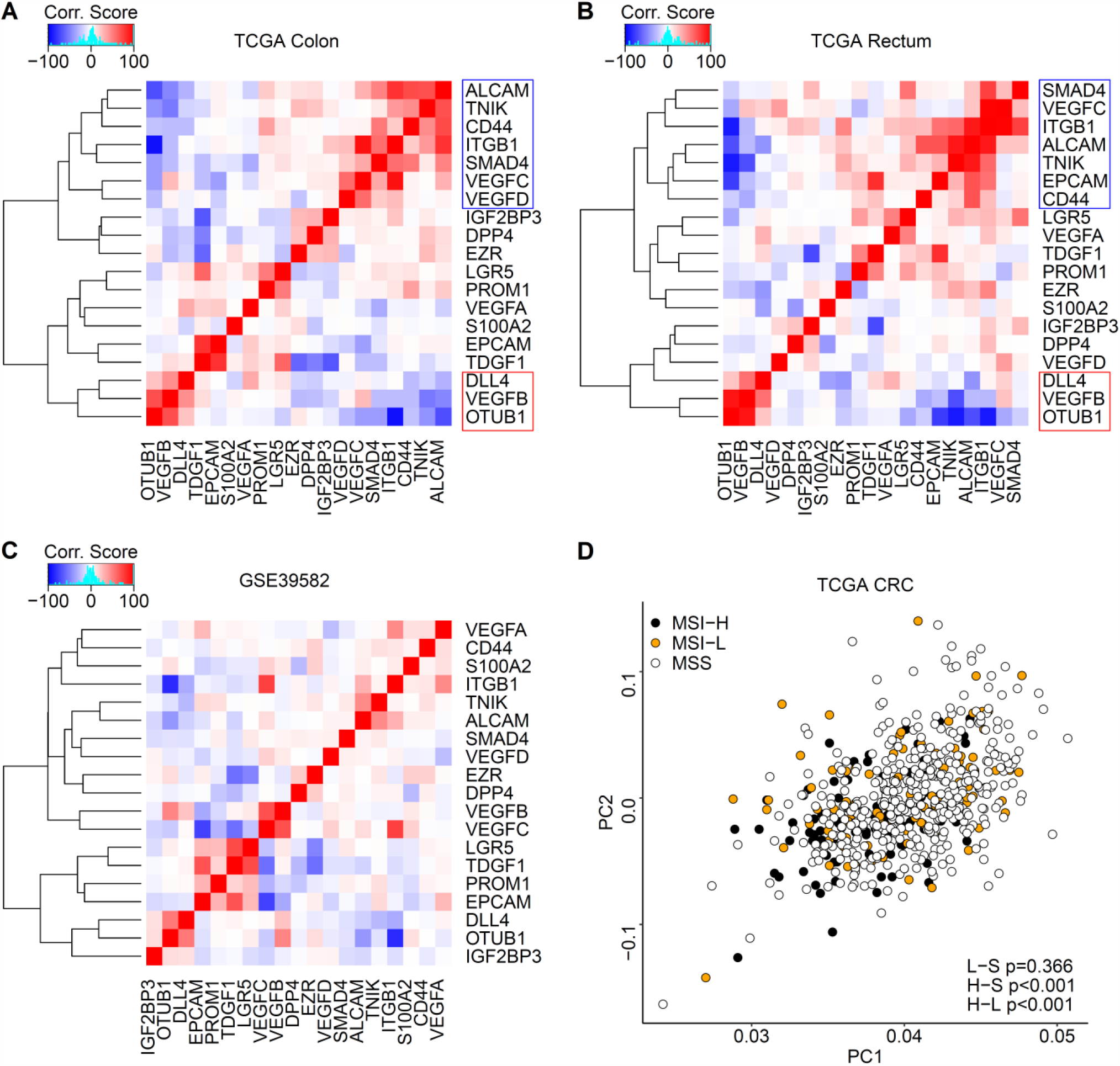
The co-expression pattern of 9 gene markers cannot be explained by the microsatellite (MS) status alone. **A and B**. A representative gene co-expression heatmap across TCGA colon cancers (**A**) highlights two gene clusters. Six of 7 genes are shared in the first (blue) cluster and all three genes are shared in the second (red) cluster with the TCGA rectum heatmap (**B**). Heatmap colors indicate the Correlation Scores between gene-pairs (see Methods). **C**. A similar heatmap built using microarray data. The 9 genes identified in the TCGA co-expression heatmaps (**A** and **B**) exhibit a similar co-expression pattern and most of them fall within the same clusters in the microarray analysis. **D**. The MS status plotted in the two-dimensional space of PC1 and PC2 for the 9 selected gene markers across TCGA CRC tumors. In the right bottom corner, MSI-L is presented as L, MSS as S, MSI-H as H and the permutation test *P* value from cluster centroid comparison (See Methods) as p. Hence, MSI-H class is significantly different from MSS and MSI-L.

Since our method employs random sampling, we repeated our clustering pipeline 100 times to verify the stability of these two gene clusters. Cluster 1’s six genes clustered together in 93 of 100 random runs using colon samples and in 44 of 100 runs using rectal samples. Cluster 2’s genes clustered together in 67 and 80 of 100 runs using colon and rectal samples, respectively. Running the same pipeline on microarray samples results in a similar pattern in the relationship among these 9 genes although not all genes within each group fall in the same cluster (Figure 2C). The heatmaps in Figure 2A, B and C are representative results of the clustering pipeline.

We then asked if the expression pattern seen in Clusters 1 and 2 can be explained by the MS status of CRC tumors. We performed PCA analysis using the 9 gene markers in all TCGA CRC samples. We examined the association between the MS classes, PC1 (explains 88.4% of the sample variations) and PC2 (explains 4.7% of the sample variations). Our results suggest that the MS status has an influence (*P* < 0.001) on the expression of these 9 genes (Figure 2D). Both datasets indicate that this influence is clearly stronger on PC1 than on PC2 (Figure S1). However, we found no clear separation of CRC samples based on their MS status (Figure 2D).

### PC2 values of the 9 gene makers divide CRC samples, particularly CMS2 and CMS4, into two subtypes

We next asked if the observed variations in the 9 gene markers are driven by CRC subtypes beyond the MS status. To answer this question, we attempted to separate CRC samples based on these gene variations. We generated heatmaps of the gene expressions across CRC samples, sorted by their PC1 values (Figures 3A and S2A) and PC2 values (Figures 3B and S2B). PC1-sorted heatmaps did not clearly separate the CRC samples. However, PC2 values identified two opposite gene expression patterns both in TCGA (Figure 3B) and microarray samples (Figure S2B). The results revealed two clear subtypes, which we named subtype 1 (S1) and subtype 2 (S2). S1 corresponds to the overexpression of Cluster 1 genes whereas S2 corresponds to the overexpression of Cluster 2 genes (Figures 3B, S2B and S3).

**Figure 3.**
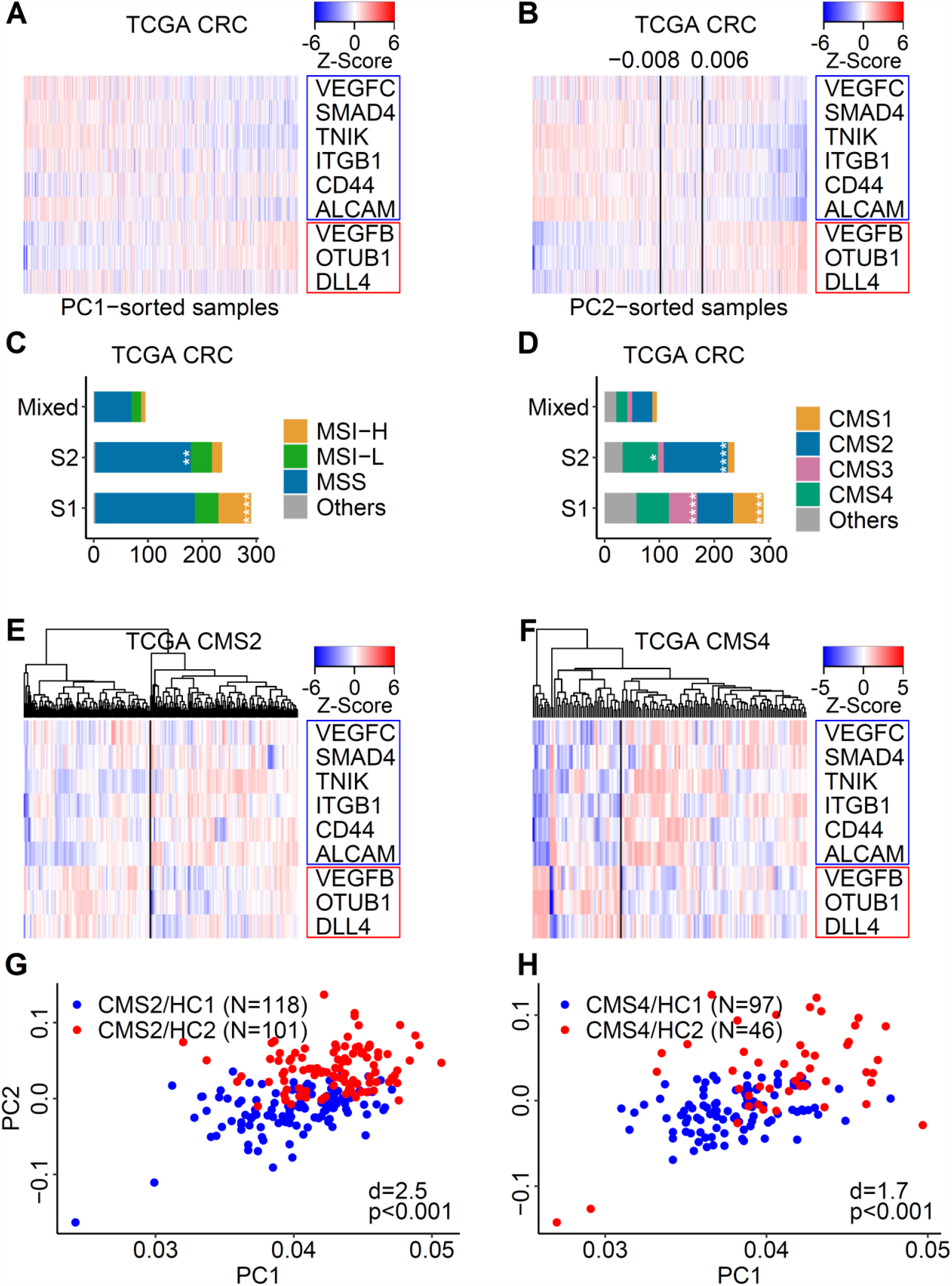
PC2 values of the 9 gene makers divide CRC samples, particularly CMS2 and CMS4, into two subtypes. **A**. A heatmap of the expression values for the 9 gene markers in TCGA CRC samples, sorted by their PC1 values. No clear subtypes are separated in this representation. **B**. A similar heatmap, sorted by samples’ PC2 values, successfully separates these samples into clear subtypes. PC2 value cutoffs, indicated in black vertical lines, separate S1, Mixed and S2 samples, respectively, from left to right. **C**. The distribution of MS classes across S1 and S2 in TCGA. Significantly enriched groups via Fisher’s exact test are indicated (*: *P*<0.05, **: *P*<0.01, ***: *P*<0.001 and ****: *P*<0.0001). **D**. The distribution of CMS classes across S1 and S2 in TCGA presented as in **C**. **E and F**. Unsupervised hierarchical clustering of TCGA CMS2 (**E**) and CMS4 (**F**) samples. Black vertical lines separate the samples of each into major clusters. **G and H**. Scatter plots of CMS2 (**G**) and CMS4 (**H**) hierarchical clusters (HC1 and HC2 obtained in **E** and **F**) across the two-dimensional space of PC1 and PC2 values. The number of samples for each hierarchical cluster, N, is shown between parentheses. Indicated in the bottom right corner: d is the Euclidian distance between HC1 and HC2 centroids after normalizing the values of each axis and p is the permutation *P* value from cluster centroid comparison (see Methods).

We then checked S1 and S2 compositions of MS and CMS classes. In TCGA, both S1 and S2 are composed of microsatellite instable-high (MSI-H), MSI-low (MSI-L) and MSS tumors. However, MSI-H is strongly enriched in S1 and MSS is strongly enriched in S2 (Figure 3C). In microarray, we found no statistically significant bias between the MS classes (Figure S2C). On the other hand, both datasets indicate that CMS1 and CMS3 mainly occur within S1 while CMS2 and CMS4 are divided between S1 and S2, with a significantly larger portion of CMS2 falling inside S2 (Figures 3D and S2D).

To verify the results of CMS relationship with S1/S2 subtypes, we performed unsupervised hierarchical clustering of samples of each CMS class separately using the 9 gene markers. Within TCGA data, CMS1 and CMS3 sample clustering did yield clusters without clear gene expression patterns (data not shown), consistent with their shortage of S2 samples. On the other hand, both CMS2 and CMS4 samples clustered into two major groups (Figures 3E, and F). We named the two groups HC1 and HC2, for hierarchical cluster 1 and 2. When we plotted HC1 and HC2 samples in the two-dimensional space of PC1 and PC2, we found them to be clearly separated, mainly by their PC2 values (Figures 3G and H). Microarray data analysis leads to the same conclusion (Figures S2E, F, G and H). These results confirm that each of CMS2 and CMS4 has at least two different subtypes.

### CMS2/S1 and CMS4/S1 samples harbor molecular features different from CMS2/S2 and CMS4/S2

We investigated the molecular differences between S1 and S2 within CMS2 and CMS4 samples. We first examined these differences using other published CRC subtyping systems: TCGA gastrointestinal tract cancer subtypes [16] and colorectal cancer intrinsic subtypes (CRIS) [29]. TCGA subtypes do not indicate differences between CMS2/S1 and CMS2/S2 (Figure S4A), but they reveal that CMS4/S1 has more genome stable tumors while CMS4/S2 has more CIN tumors (Figure S4B). This subtyping is not available for microarray data. Our analysis of CRIS subtypes indicates that CRIS-C, with elevated EGFR signaling, is enriched in CMS2/S2 compared to CMS2/S1 (Figures S4C and E). Meanwhile, CRIS-B, characterized by TGF-β activation, is more prevalent in CMS4/S1 compared to CMS4/S2 (Figures S4D and F).

Regarding the methylation status, we found that CMS2 tumors, whether S1 or S2, rarely harbor highly methylated (CIMP-H in TCGA or CIMP in microarray) tumors (Figures 4A and S4G). CMS2 and CMS4 samples classified as S1 exhibit a clear enrichment of lowly methylated (CIMP-L) tumors (Figures 4A and B), a feature assessed only in TCGA data. In addition, CIMP-H/CIMP tumors are more prevalent in CMS4/S1 compared to CMS4/S2 (Figures 4B and S4H).

**Figure 4.**
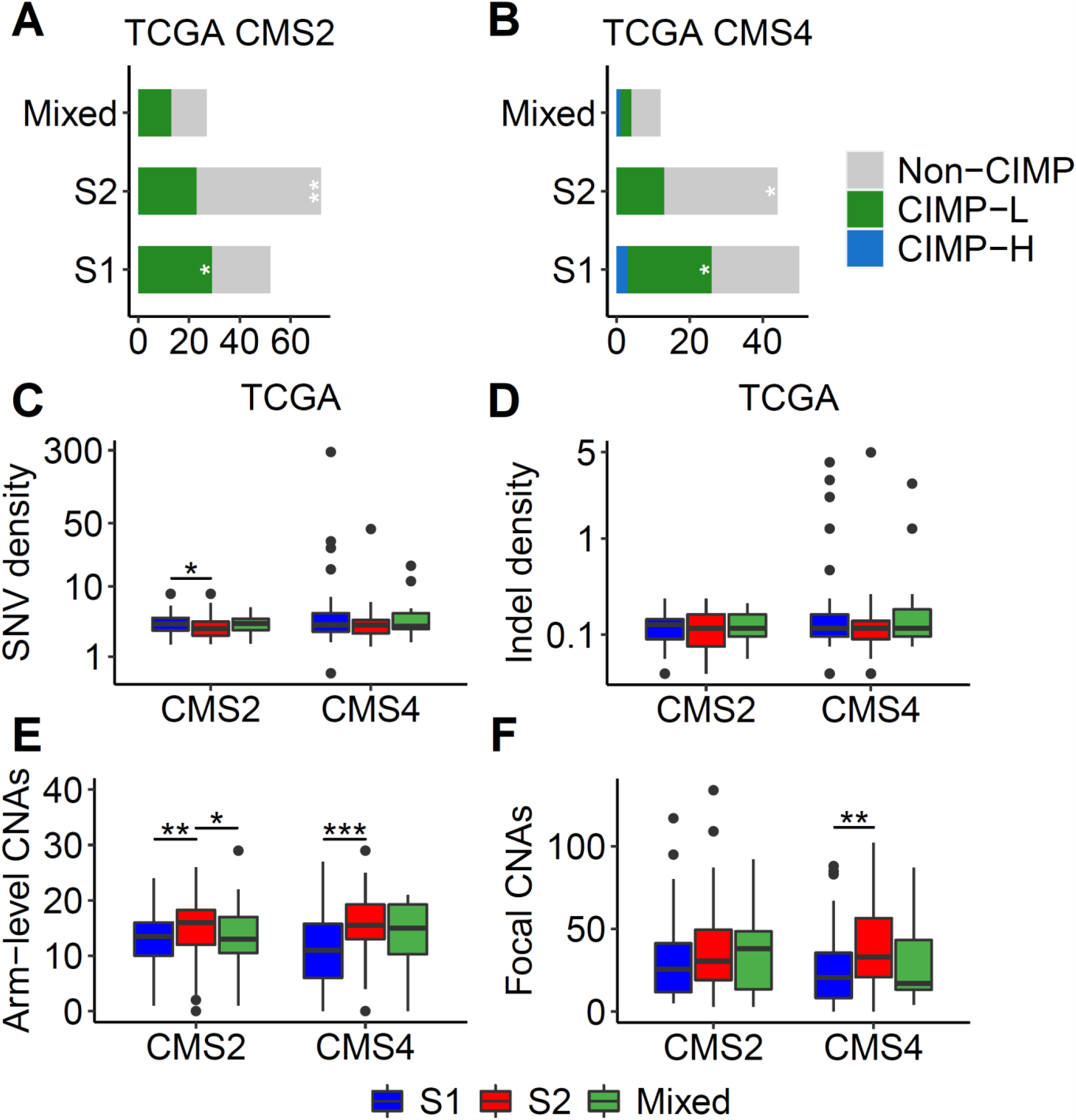
CMS2/S1 and CMS4/S1 samples harbor molecular features different from CMS2/S2 and CMS4/S2. **A and B**. S1 and S2 compositions based on their methylation status in TCGA CMS2 (**A**) and CMS4 (**B**) samples. CIMP: CpG island methylated phenotype, Non-CIMP: non-methylated, CIMP-L: CIMP-low and CIMP-H: CIMP-high. **C**. The distribution of somatic single nucleotide variation (SNV) density in TCGA CMS2 and CMS4 tumors. Significant differences within the same CMS class via Wilcoxon signed-rank test are indicated. **D, E and F**. The distribution of small somatic indel density (**D**), arm-level (**E**) and focal (**F**) somatic copy number alterations (CNAs) in TCGA CMS2 and CMS4 tumors presented as in **C**.

Finally, we looked at the mutation density, arm-level and focal copy number alterations (CNAs), which are available only for TCGA data. Whether in CMS2 or CMS4, S1 harbors higher single nucleotide variation and indel densities (Figures 4C and D) while S2 harbors higher arm-level and focal CNAs (Figures 4E and F). Our collective analysis concludes that S1 and S2 of CMS2 and CMS4 have different molecular features.

### A strong enrichment of six cancer hallmarks highlights the S1 subtype across all CMSs

Next, we asked whether the S1 and S2 difference is limited to the expression of the 9 gene markers or is extended to the expression of more genes and pathways. We performed S1 versus S2 enrichment analysis for all cancer hallmarks [19] across all CMSs. We used four enrichment methods in this part: GSEA, ssGSEA [30], GSVA [31] and Fisher’s exact test of differentially expressed genes overlapping each hallmark. The analysis of TCGA dataset suggests that six hallmarks (androgen response, protein secretion, mitotic spindle, UV response down, PI3K and TGF-β signaling) are strongly enriched in S1 across all CMS classes (Figure 5). Similar results were observed using the microarray dataset (Figure S5). Indeed, TGF-β enrichment in S1 is consistent with the CRIS-B prevalence within CMS4/S1 observed in our previous analysis.

**Figure 5.**
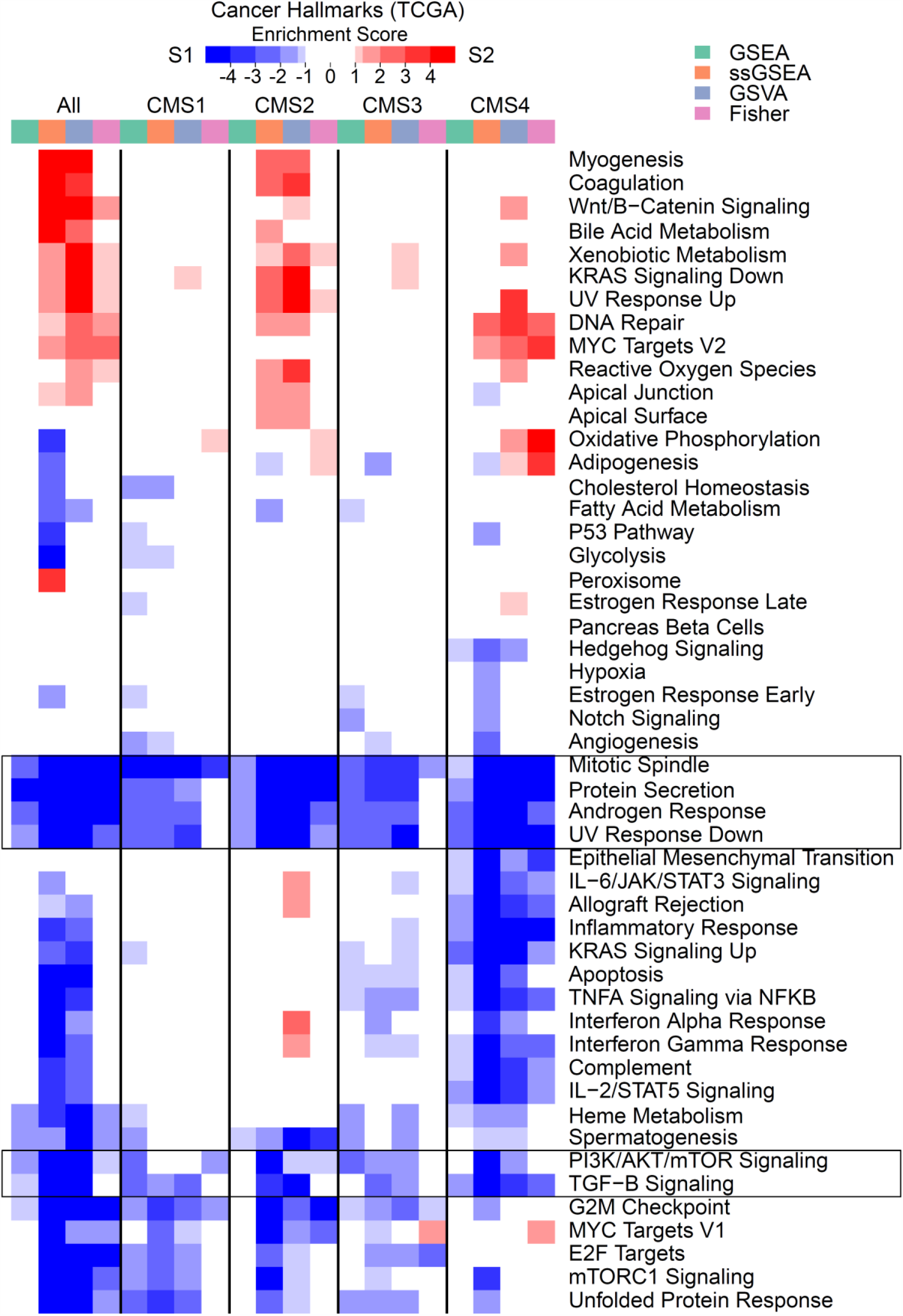
A strong enrichment of six cancer hallmarks highlights the S1 subtype across all CMSs. A heatmap of cancer hallmark pathway enrichment (S1 versus S2) within CMS classes of TCGA samples using four methods. The six pathways marked by black rectangles are clearly activated in S1 samples across all CMSs. The Enrichment Score is −log_10_(*P*). A negative sign is assigned for S1 enrichment. Nominal *P* values were used for GSEA and two-sided t-test *P* values were used for ssGSEA and GSVA methods. In Fisher’s method, the significance of the overlap between each pathway genes and differentially expressed genes between S1 and S2 at two-tailed t-test *P*=.0001 was used.

We observed that in TCGA data, many of the immune response pathways (including IL-6 and interferon alpha and gamma) are activated in S1 of CMS4 and in S2 of CMS2. These signals still exist but are not significant in the microarray samples.

### The tumor microenvironment is activated in S2 samples of CMS2 and S1 samples of CMS4

Both TCGA and microarray indicate a depletion of the epithelial signature in S2, particularly within CMS2. Furthermore, both datasets show that S2 samples within CMS2 are characterized by the activation of the tumor microenvironment. Indeed, almost all immune cells but T-helper and T central memory (Tcm) cells are activated in S2 of CMS2 (Figures 6A and S6A). Other stromal components, both the invasive front and leukocyte signatures, are also enriched in the same samples (Figures 6B and S6B). CMS4 samples, however, differ. Specifically, in TCGA, we observed a strong immune activation (Figure 6A) and a likely stromal activation (Figure 6B) in S1 of CMS4. We found no consistent association across the two datasets between the OS and any of the microenvironment components activated in S1 or S2 of CMS2 or CMS4 (Figures 6 and S6). These results reiterate the molecular differences between S1 and S2 within CMS2 and CMS4 indicated previously.

**Figure 6.**
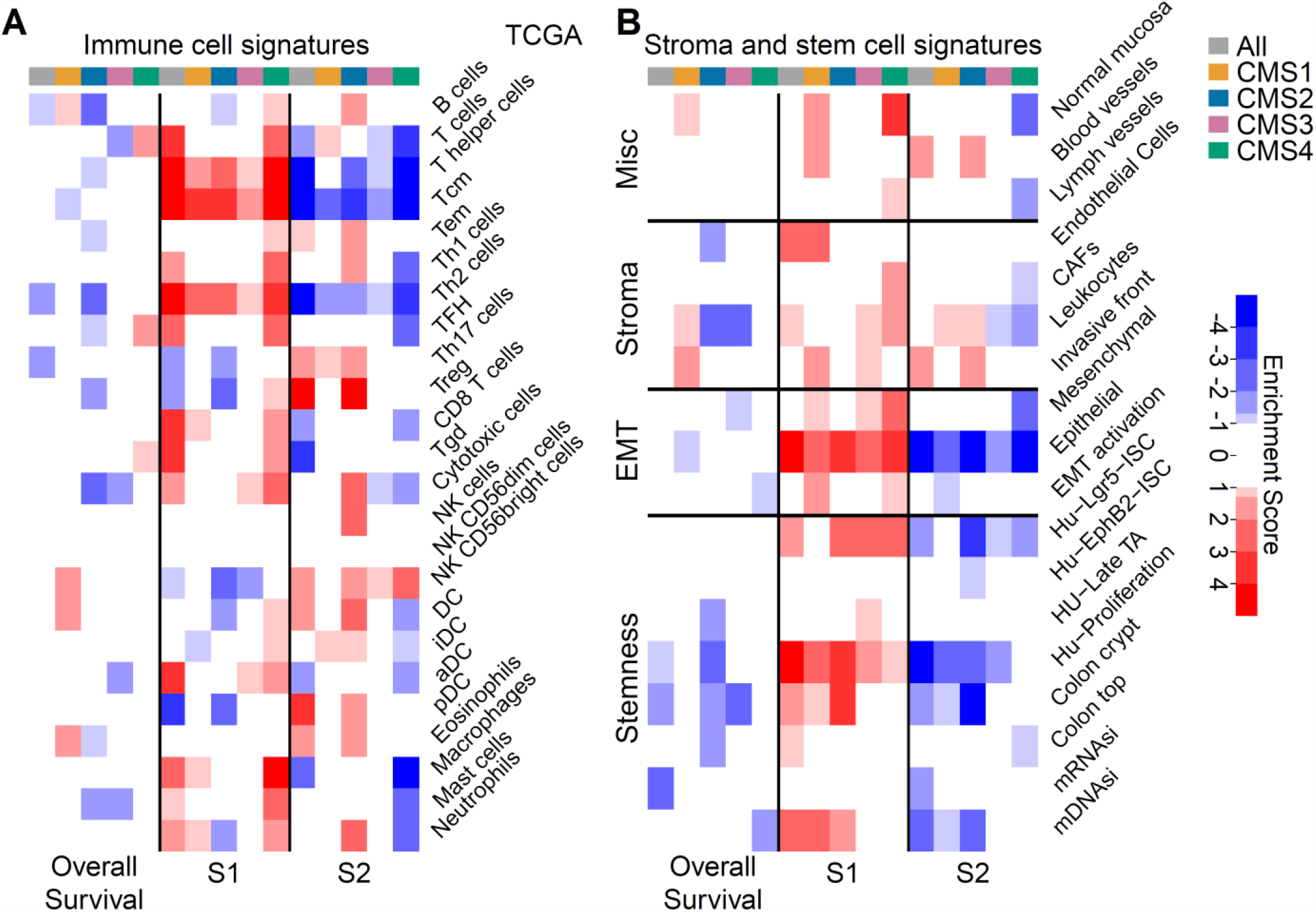
The tumor microenvironment is activated in S2 samples of CMS2 and S1 samples of CMS4. **A**. A heatmap of the relationship between immune cell signatures via ssGSEA score (see Methods), the overall survival (OS), S1 and S2 in TCGA data. OS is better (blue) or worse (red), according to Kaplan-Meier log-ranked test *P* value, for samples bearing the indicated signature. A signature is depleted (blue) or enriched (red) in each group of S1 or S2 in a CMS class indicated by the column labels. **B**. Also using TCGA, a heatmap of the relationship between tumor microenvironment components, OS, S1 and S2 presented as in **A**.

### S2 is associated with lymphatic invasion across CMSs and more frequent metastasis in CMS2

At the end, we decided to examine the clinical outcome of S1 and S2. Within TCGA, the invasive signature of S2 in CMS2, observed previously, is buttressed by its strong association with lymphatic invasion events in all CRCs (*P* = .0002; Figure S7A) and in CMS2 (*P* = .012; Figure 7A). This pattern still holds but not as significant within CMS4 (Figure 7B). Unfortunately, the lymphatic invasion assessment is not available for the microarray samples to validate these observations.

**Figure 7.**
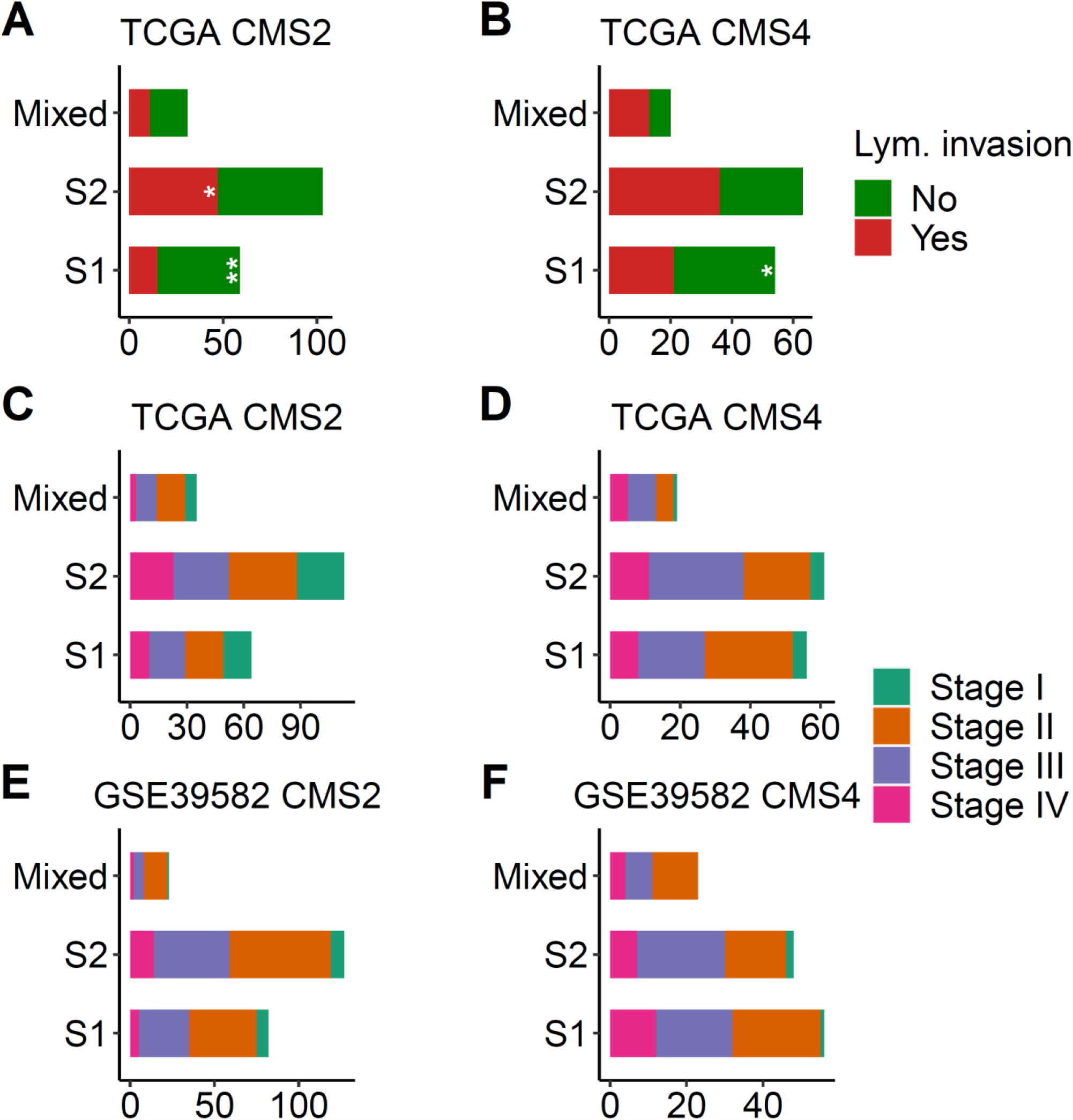
S2 is associated with lymphatic invasion across CMSs and more frequent metastasis in CMS2. **A and B**. The distribution of lymphatic invasion events across S1 and S2 of TCGA CMS2 (**A**) and CMS4 (**B**) cases. **C, D, E and F**. The distribution of pathological stages across S1 and S2 of TCGA (**C** and **D**) and microarray (**E** and **F**) CMS2 and CMS4 cases.

For the pathological stage, the only consistent pattern in both datasets indicates that Stage IV tumors are more frequent in CMS2/S2 (20.4 and 11% in TCGA and microarray, respectively) compared to CMS2/S1 (15.6 and 6.1%, respectively) (Figures 7C and E). We found no consistent association between the pathological stage and S1 or S2 within CMS4 (Figures 7D and F) or across all CRCs (Figures S7B and C).

Finally, TCGA data indicate that CMS2/S2 patients have a relatively worse OS (Figure S7D). This observation is not confirmed by the microarray data (Figure S7F). Conversely, the microarray data, but not TCGA, show that CMS4/S1 patients have a relatively worse OS than CMS4/S2 patients (Figures S7E and G). Overall, we conclude that S2 is more invasive and can pose a higher risk of metastasis for CMS2 patients.

## Discussion

In this study, we used previously reported metastasis and CSC markers specific to CRC. A subset of the starting markers serves as a simplified signature for CRC subtypes that exist within CMS2 and CMS4 samples. The first subtype, S1, corresponds to the overexpression of six gene markers: *ALCAM, TNIK, CD44, ITGB1, SMAD4* and *VEGFC*. The second subtype, S2, corresponds to the overexpression of three markers: *VEGFB, OTUB1* and *DLL4*.

In a recent study, mouse xenografts were used to identify CRC intrinsic molecular subtypes [29]. The study found that most of CMS1 and CMS3 samples fall into one CRC intrinsic subtype, CRIS-A, which is mostly MSI. To the contrary, CMS4 samples are distributed between five intrinsic subtypes and most CMS2 samples fall into three intrinsic subtypes. Our simple 9-gene markers lead to subtypes (S1 and S2) that indicate a similar observation regarding CMS heterogeneity. Our study shows that CMS1 and CMS3 are strongly biased towards S1 whereas both CMS2 and CMS4 have significant portions distributed between S1 and S2 (Figures 3F and H).

We demonstrate that the methylation status, mutation density and CNA level of CMS2 and CMS4 are important distinctions in their heterogeneity at the molecular level. In alignment with these results, a recent gastrointestinal tract pan-cancer study suggests that the methylation status may have an important role in the evolutionary process of CRC CIN tumors [16], which are mostly CMS2 and CMS4.

Our analysis indicates that six cancer hallmarks are strongly enriched in S1 samples. Further analysis reveals that S2 of CMS2 is associated with markers of elevated immune response, stromal invasion and tumor invasive front. These CMS2/S2 molecular signatures are accompanied by a significant increase in lymphatic invasion and a higher frequency of metastasis.

We further investigated the clinical implications of the S1/S2 subtypes within CMS2 or CMS4 via OS (Figure S7). We could not find statistically significant and consistent results in this analysis. For example, TCGA data suggest that S2 is linked to worse OS within CMS2 (*P* = .1), in agreement with its invasiveness, but there was no detected OS difference in microarray data. This inconsistency might be related to the insufficient sample size to perform a conclusive survival analysis.

Our study does not provide an insight about the mechanistic means of S2 invasiveness and metastasis in CMS2. CMS4 is believed to maintain its invasiveness through the mesenchymal transformation, which is linked to the activation of the tumor microenvironment. Based on our molecular analysis, S2 features differ between CMS2 and CMS4 (Figures 6 and S6). This raises the possibility that S2 molecular features might be required for CMS2 invasiveness and metastasis but not for CMS4. If true, then CMS2 might acquire its invasive phenotype through a non-mesenchymal path. The high copy number alteration, which is a feature of CMS2/S2, and has an established link to worse clinical outcome in multiple cancers, including CRC [32], is one possible path.

In conclusion, our study shows, based on the two largest public CRC gene expression datasets, that CMS2 and CMS4 are composed of two molecularly distinct subtypes: S1 and S2. The S2 subtype has multiple molecular signatures of invasiveness and a potentially worse clinical outcome within CMS2 cases.

## Acknowledgement

This research project has been funded by King Abdulaziz City for Science and Technology.

